# Mitochondrial protein heterogeneity stems from the stochastic nature of co-translational protein targeting in cell senescence

**DOI:** 10.1101/2023.12.03.569753

**Authors:** Abdul Haseeb Khan, Rutvik J. Patel, Matheus P. Viana, Susanne M. Rafelski, Aidan I. Brown, Brian M. Zid, Tatsuhisa Tsuboi

**Affiliations:** Institute of Biopharmaceutical and Health Engineering, Tsinghua Shenzhen International Graduate School, Shenzhen 518055, China; Department of Physics, Toronto Metropolitan University, Toronto, M5B 2K3, Canada; Allen Institute for Cell Science, Seattle, WA 98109, USA; Department of Chemistry and Biochemistry, University of California San Diego, La Jolla, CA 92023, USA; Tsinghua-SIGS & Jilin Fuyuan Guan Food Group Joint Research Center, Tsinghua Shenzhen International Graduate School, Shenzhen 518055, China

**Keywords:** mitochondria, noise, cell senescence, translation, mRNA localization

## Abstract

A decline in mitochondrial function is a hallmark of aging and neurodegenerative diseases. It has been proposed that changes in mitochondrial morphology, including fragmentation of the tubular mitochondrial network, can lead to mitochondrial dysfunction, yet the mechanism of this loss of function is unclear. Most proteins contained within mitochondria are nuclear-encoded and must be properly targeted to the mitochondria. Here, we report that sustained mRNA localization and co-translational protein delivery leads to a heterogeneous protein distribution across fragmented mitochondria. We find that age-induced mitochondrial fragmentation drives a substantial increase in protein expression noise across fragments. Using a translational kinetic and molecular diffusion model, we find that protein expression noise is explained by the nature of stochastic compartmentalization and that co-translational protein delivery is the main contributor to increased heterogeneity. We observed that cells primarily reduce the variability in protein distribution by utilizing mitochondrial fission-fusion processes rather than relying on the mitophagy pathway. Furthermore, we are able to reduce the heterogeneity of the protein distribution by inhibiting co-translational protein targeting. This research lays the framework for a better understanding of the detrimental impact of mitochondrial fragmentation on the physiology of cells in aging and disease.

## INTRODUCTION

Mitochondria are hubs for metabolites and energy generation and have been deemed important for age-related processes and diseases, including cancer and neurodegeneration ^1–3^. The loss of mitochondrial function in these pathological phenotypes is tied to cell-to-cell variability in gene expression caused by the amount and functional mitochondrial heterogeneity within a cell ^4–6^. Two different aspects of mitochondrial heterogeneity of a single cell are often discussed: genetic heterogeneity and functional heterogeneity ^7^. Genetic heterogeneity is based on the distribution of mtDNA and varying rates of mtDNA gene transcription and translation, which increase the heterogeneity of protein levels for each fragment ^8^, known as noise ^9^. On the other hand, definitions of functional heterogeneity are more diverse, involving cristae structure ^10,11^, membrane potential ^12,13^, and pH ^14,15^ distributions in mitochondria. Both genetic and functional heterogeneities are influenced by mitochondrial morphology, especially by fragmentation. The sustained fragmentation of mitochondria precludes the exchange of mitochondrial contents among mitochondria will lead to non-optimal protein stoichiometry of mitochondrial complexes, disruption of normal protein homeostasis, and loss of functional mitochondria ^16–23^. While fragmented mitochondria are suggested to be degraded by the mitochondria quality control pathway through mitophagy ^24^, hyper-fragmented mitochondria, which is also a phenotypic feature of diseases, require more dynamic regulatory mechanisms to eliminate the potentially harmful mitochondrial state from the cells such as mitochondrial fission-fusion reactions and tubular structure formation ^25^.

Mitochondrial proteins are encoded mainly in the nuclear genome, which are then imported to mitochondria from the cytoplasm ^26^. A large fraction of mitochondrial protein-coding mRNA is localized to the mitochondrial outer membrane and co-translationally imports proteins into mitochondria ^27–33^. The mechanism of mRNA localization is based on both the 3′ UTR and coding regions, primarily through mitochondrial targeting sequences (MTSs) ^28,31–35^. While there is an advancing understanding of mitochondrial mRNA localization mechanisms, the impact of co-translational protein targeting on mitochondrial function under different mitochondrial morphologies is poorly understood. Furthermore, mtDNA only encodes a small number of genes, eight protein-coding genes in *Saccharomyces cerevisiae* ^36^, raising the question of whether the distribution of nuclear-encoded proteins, which are imported into mitochondria from the cytoplasm, causes mitochondrial dysfunction. Here, we report that co-translational protein delivery and mitochondrial fragmentation together lead to heterogeneity in protein distribution to mitochondria in aged cells. Our work shows that mitochondrial fragmentation affects protein targeting and results in higher protein heterogeneity in each mitochondrial fragment. This heterogeneity is normally homogenized by the mitochondrial fission-fusion processes, not the mitophagy pathway. By inhibiting the co-translational protein delivery, we successfully reduced protein heterogeneity. We propose that age-associated fragmentation of mitochondria induces an increased heterogeneity of mitochondrial proteins, ultimately leading to accelerated cell senescence.

## Results

### Aging increases heterogeneity in mitochondrial protein distribution in each mitochondrion

Nuclear-encoded mitochondrial protein import is essential for the proper functioning of mitochondria. Disruption of this process has been linked to a number of diseases ^37^. The core of the TIM23 complex, composed of Tim23p, Tim50p, and Tim17p, forms a translocase of the inner mitochondrial membrane that is important for importing a large number of matrix proteins ^38^. To analyze the effect of aging on the distribution of mitochondrial proteins in yeast, we adopted mother-enrichment-program (MEP) yeast strains ^39^, with which we can continuously observe mother cells by inhibiting a cell cycle progression of the daughter cell, and then determined the protein expression levels of Tim50p and Tim23p by quantitative microscopic techniques. We analyzed cells at 0 divisions (0 hours) and ∼16 divisions (24 hours) following the induction of MEP (Figure 1A, Supplementary Figure 1). Consistent with prior research ^23^, the number of fragments increased with aging (24 hours) (Figure 1B). To evaluate the mitochondrial fragmentation in each cell, we calculated the proportion that each mitochondrion in a cell contributes to the total mitochondrial volume at the single-cell level. For a cell with a single, fully tubular mitochondrion, this portion would be 1; whereas, for a cell with many highly fragmented mitochondria, the portion for each mitochondrion would be much smaller than 1, as the volume of each fragment only contributes a small fraction of the total mitochondrial volume. At 24 hours, when there are many fragments per cell, we see that the typical portion of mitochondria for each mitochondrion is small, while a much larger portion for each mitochondrion was observed at 0 hours (Figure 1C). Imbalances in protein stoichiometry can lead to the loss of protein complex function and even protein aggregation-driving proteotoxic stress ^40^. As each fragmented mitochondria can act as an independent fundamental unit, we sought to quantify the heterogeneity of Tim50 and Tim23 proteins across mitochondrial fragments. We observed greater heterogeneity in the Tim50/Tim23 relative protein expression in each mitochondrial fragment (Figure 1D) and confirmed the significant increase in the noise (coefficient of variation) level with increased age (Figure 1E). We hypothesized that the observed age-induced changes in mitochondrial morphology may contribute to this increased noise level, so we analyzed the relationship between the protein expression level and mitochondrial fragment size. The increased number of smaller mitochondrial fragments during aging was linked to the increased heterogeneity of the Tim50/Tim23 relative protein levels (Figure 1F, G). Interestingly, the overall protein expression of the fragments remained constant, as the linear regression line of Tim50/Tim23 relative protein level in Figure 1F stayed the same over the fragments. These results provide a de novo observation that aging increases heterogeneity in the distribution of mitochondrial protein at the overall fragment level.

**Figure 1.**
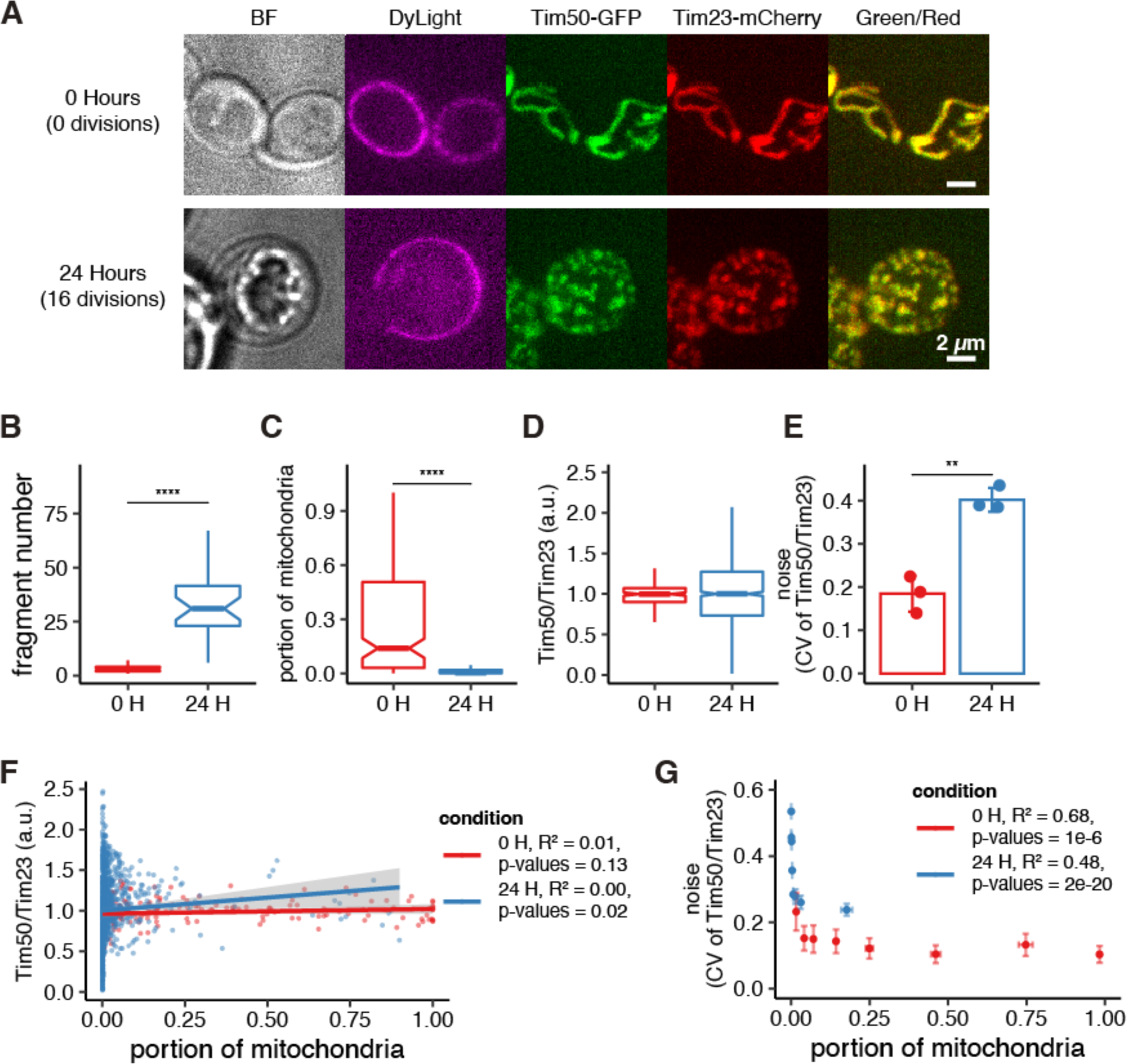
Aging increases the heterogeneity of mitochondrial protein distribution. (A) Protein distribution to each mitochondrial fragment in a young cell (top) and an aged cell (bottom). The cells of mother enrichment program strains marked with DyLight were analyzed at 0 hours and 24 hours after the addition of estradiol. The fluorescent of Tim50-GFP and Tim23-mCherry were visualized by a spinning disc confocal microscope. BF: Bright field (center focal plane), DyLight, Tim50-GFP, Tim23-mCherry: z-projected images of each channel (Far-Red, GFP, RFP). Green/Red: merge of GFP and RFP channels. Scale bar, 2 µm. (B) The number of mitochondrial fragments per cell of the cells at 0 hours and 24 hours cells in A (n>50). Statistical significance was assessed by the Mann–Whitney U-test (**** P < 0.0001). (C) The size of each mitochondrion was normalized as a portion of mitochondria per cell based on the sum of Tim23-mCherry intensity (n>50). Statistical significance was assessed by the Mann–Whitney U-test (**** P < 0.0001). (D) The ratio distribution for the protein expression level of Tim50-GFP and Tim23-mCherry for each mitochondria fragment. The median was normalized to 1. (E) The coefficient of variation (CV) or noise of data in (D). The results represent the mean ± standard deviation of three independent experiments. Statistical significance was evaluated by Student t-test (** P < 0.01) (F) The ratio distribution for the protein expression level of Tim50-GFP and Tim23-mCherry for each mitochondria fragment in (D) along with the portion of mitochondria. Each dot represents an independent mitochondrial fragment. The gray region surrounding the trend lines represents each line’s 95% confidence interval. (G) The noise of (F) along with the portion of mitochondria. Each noise data point was calculated by dividing the population of a portion of mitochondria into eight groups along the portion of mitochondria. Error bars represent SEM of three independent experiments. For (F, G), R-squared values and p-values for the F statistic hypothesis test are shown on the side.

### Stochastic nature of co-translational proteins targeting increased protein heterogeneity

Our observation suggests that small portion sizes of mitochondria cause high heterogeneity in protein expression, leading to increased protein expression heterogeneity in aged cells with predominantly small mitochondria. Most mitochondrial proteins are imported from the cytoplasm; however, the mRNA number in a single cell is limited to 5-10 molecules or fewer for most genes ^40–42^. Since the limited number of reactant molecules can impact the rate of a chemical reaction in general, we sought the quantitative relationship of how mRNA numbers and co-translational protein targeting to mitochondria affect mitochondrial protein heterogeneity. To explore this, we analyzed the single molecule mRNA movement of *TIM50* mRNA with a relationship to mitochondria in live cells. We visualized mitochondria using mCherry linked to the mitochondrial matrix marker, Su9, while mRNAs were labeled with the MS2-MCP system. We reconstructed and analyzed the 3D mRNA movement trajectory using Mitograph V2.0 and Trackmate (ImageJ Plugin) (Figure 2A) ^33^. We found that *TIM50* mRNAs localized to mitochondria (blue dots) showed restricted and slower movement as compared to freely moving mRNAs (red dots) (Figure 2B). Moreover, the distance traveled by mRNA localized to the mitochondrial surface is also reduced compared to freely moving mRNAs (Figure 2C). This leads mRNAs to remain localized to small mitochondrial regions for long periods. Collectively, these observations indicate that mitochondrial localization restricts the movement of *TIM50* mRNAs. Considering the half-life and abundance ratio of mRNA and protein, single mRNAs can produce tens to hundreds of proteins over an mRNA lifetime ^43^. This suggests that the limited number of mRNA largely provides translated protein into a fixed number of mitochondria when the mitochondria become morphologically fragmented.

**Figure 2.**
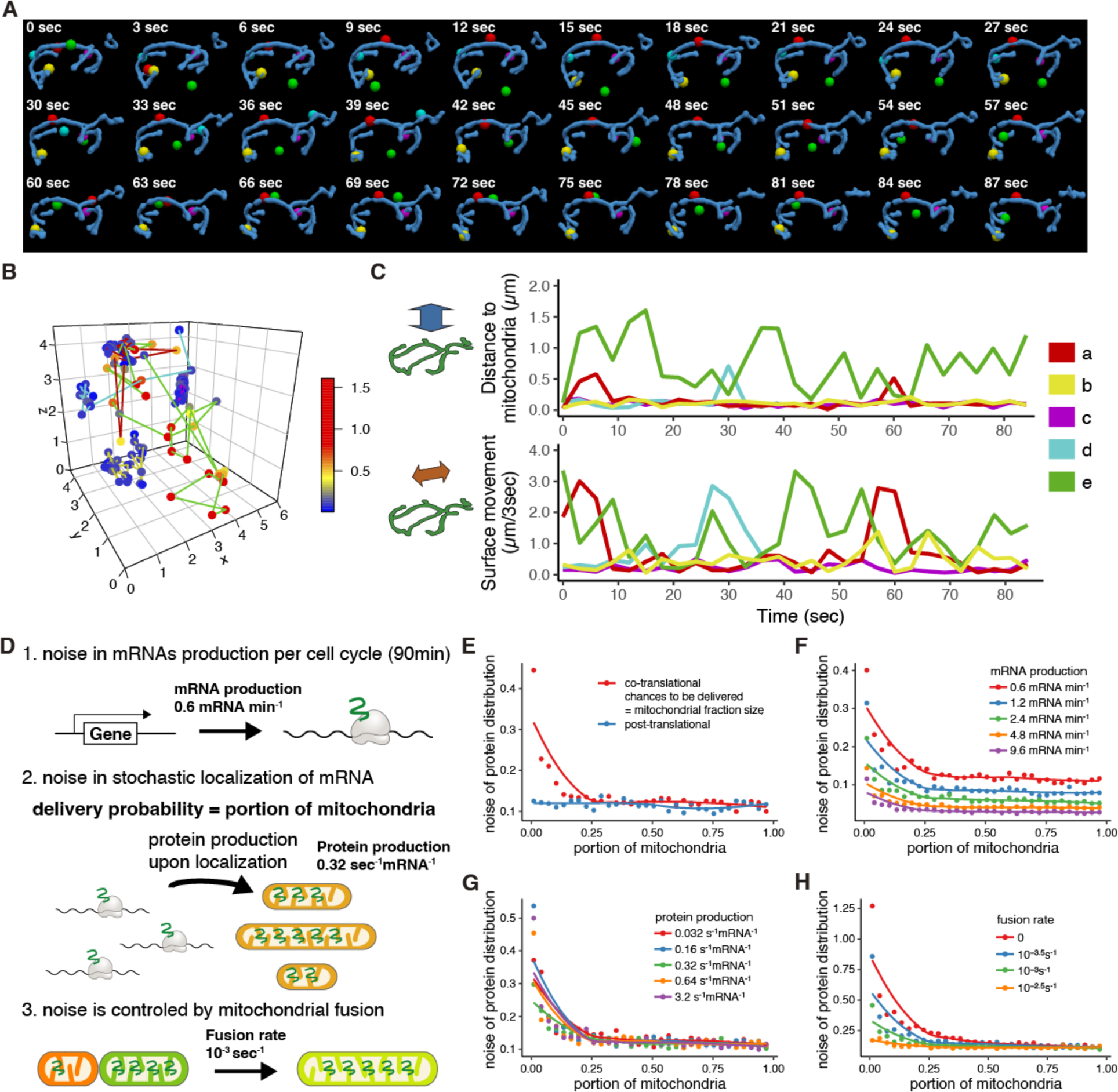
Stochastic nature of co-translational protein targeting increases the protein heterogeneity at the single mitochondrion. (A) Reconstruction of mitochondrial and mRNA positional information. Blue: mitochondria. The color particle and lines correspond to the same mRNAs (A-C). (B) The 3D trajectory of mRNA movement. The colors of mRNA positions indicate the distance from the mitochondrial surface. Distance units for 3D trajectory and color bar indicating distance from mitochondria are in units of μm. (C) The 2D trajectory of mRNA movements regarding the distance to mitochondria (µm) and the horizontal movement with respect to the surface (surface movement) in 3 seconds (µm/3sec). (D) A stochastic model description in which mitochondrial size and co-translational protein targeting with mitochondrial fission-fusion coordinate protein distribution using translational kinetic and molecular diffusion. (E-H) Relationship between the noise of protein distribution and mitochondrial size of different protein import mechanisms (E), mRNA production rates (F), protein production rates (G), and fission-fusion rates (H) in the stochastic model. The trend line was calculated using a Loess regression.

To explore the underlying causes of protein expression noise, we apply mathematical modeling (Figure 2D). With a *TIM50* mRNA copy number of six ^33^ and a lifetime of 10 minutes ^44,45^, there will be a mean of 54 mRNA produced in a 90 minute generation, with a corresponding noise of 0.14. If the number of proteins produced is proportional to mRNA produced, then protein numbers will have a similar noise due to mRNA production stochasticity of 0.14, similar to the minimum in Fig. 1G, setting a size-independent noise floor. In contrast, with protein delivery proportional to mitochondrial size, the noise will be inversely proportional to the square root of the mitochondrial portion size, such that the noise will decrease with increasing mitochondrial size with decreasing slope for larger mitochondria, similar to the trend observed in Figure 1G. With post-translational protein delivery, there are a large number of independent delivery events, leading the mitochondrial size-dependent noise from protein delivery to specific mitochondria to be relatively low. In contrast, with co-translational protein delivery, there are fewer protein delivery events, and the noise from protein delivery is relatively high.

The stochasticity from mRNA production and protein delivery are combined in a stochastic simulation (Figure 2D). Post-translational protein delivery leads to noise that is independent of size and slightly above mRNA production noise of 0.14 (Figure 2E). In contrast, co-translational protein delivery leads to a size-dependent noise, with greater noise for small mitochondria (Figure 2E) that can be increased by decreasing the mRNA production rate (Figure 2F). It is interesting to note that the rate of protein production by each mRNA does not affect the noise (Figure 2G). Mitochondrial fusion facilitates protein spread between mitochondria, such that with co-translational protein delivery, the mitochondrial size dependence of the noise is enhanced by a low mitochondrial fusion rate. Increasing the fusion rate compresses the size range over which noise increases to the smallest mitochondria, and at the highest mitochondrial fusion rates, the size dependence of noise is eliminated, with only noise due to mRNA production remaining (Figure 2H).

### Mitochondrial fusion dynamics maintain the homogeneous protein distribution

Mitochondrial morphological changes are dictated by the balance of fission-fusion reactions and mitophagy pathways maintained by regulatory proteins ^3,46^. To investigate whether mitochondrial fusion dynamics regulate the homogeneity of mitochondrial protein distribution, we used the fusion mutant, *fzo1*Δ. As expected, cells lacking Fzo1 exhibited fragmented mitochondrial morphology and a significant increase in the noise level of relative Tim50/Tim23 protein expression (Figure 3A, B). Moreover, consistent with the above modeling (Figure 2H), the noise level decreased substantially with an increase in the portion size of mitochondria in *fzo1*Δ cells (Figure 3C). We next used the mitophagy deficient strain, *atg32*Δ, to examine the role of mitophagy in protein heterogeneity. In contrast to *fzo1*Δ cells, we found that *atg32*Δ cells showed no significant increase in mitochondrial fragmentation (Figure 3D) or the noise level of relative Tim50/Tim23 protein expression (Figure 3E). Heterogeneity decreased with increasing mitochondrial portion size for *atg32*Δ cells, similar to the WT (Figure 3F). To further examine if these regulatory relationships can be applied to general mitochondrial protein homeostasis, we tested two other mitochondrial proteins that are not the components of an identical protein complex. We co-expressed Tim50-GFP and Su9-mCherry, a subunit 9 of mitochondrial ATPase of *Neurospora crassa* ^47^ conjugated with mCherry and localized to the mitochondrial matrix in yeast strains. We observed fragmented mitochondrial morphology and significantly increased noise level in Tim50/Su9 relative protein expression in the *fzo1*Δ cells, compared to WT (Figure 3G, H). Furthermore, Tim50/Su9 heterogeneity remains consistently high in fusion mutant cells, even for large mitochondrial portions (Figure 3I). This is yet very different from the exponential relationship which was seen in the noise of Tim50/Tim23 (Figure 3C). This suggests that when proteins do not form a complex, the homogeneity of proteins can easily be dysregulated. We observed a similar expression phenotype for relative Tim50/Su9 and Tim50/Tim23 proteins in atg32Δ cells (Figure 3J-L). All the above evidence suggests that fission-fusion dynamics are responsible for regulating heterogeneity in mitochondrial protein expression, and mitophagy may only have a minor influence on mitochondrial protein heterogeneity. These observations are not restricted to a specific protein complex as Tim50/Tim23 and Tim50/Su9 relative protein expression levels show similar noise level phenotypes as mitochondrial fusion dynamics increase.

**Figure 3.**
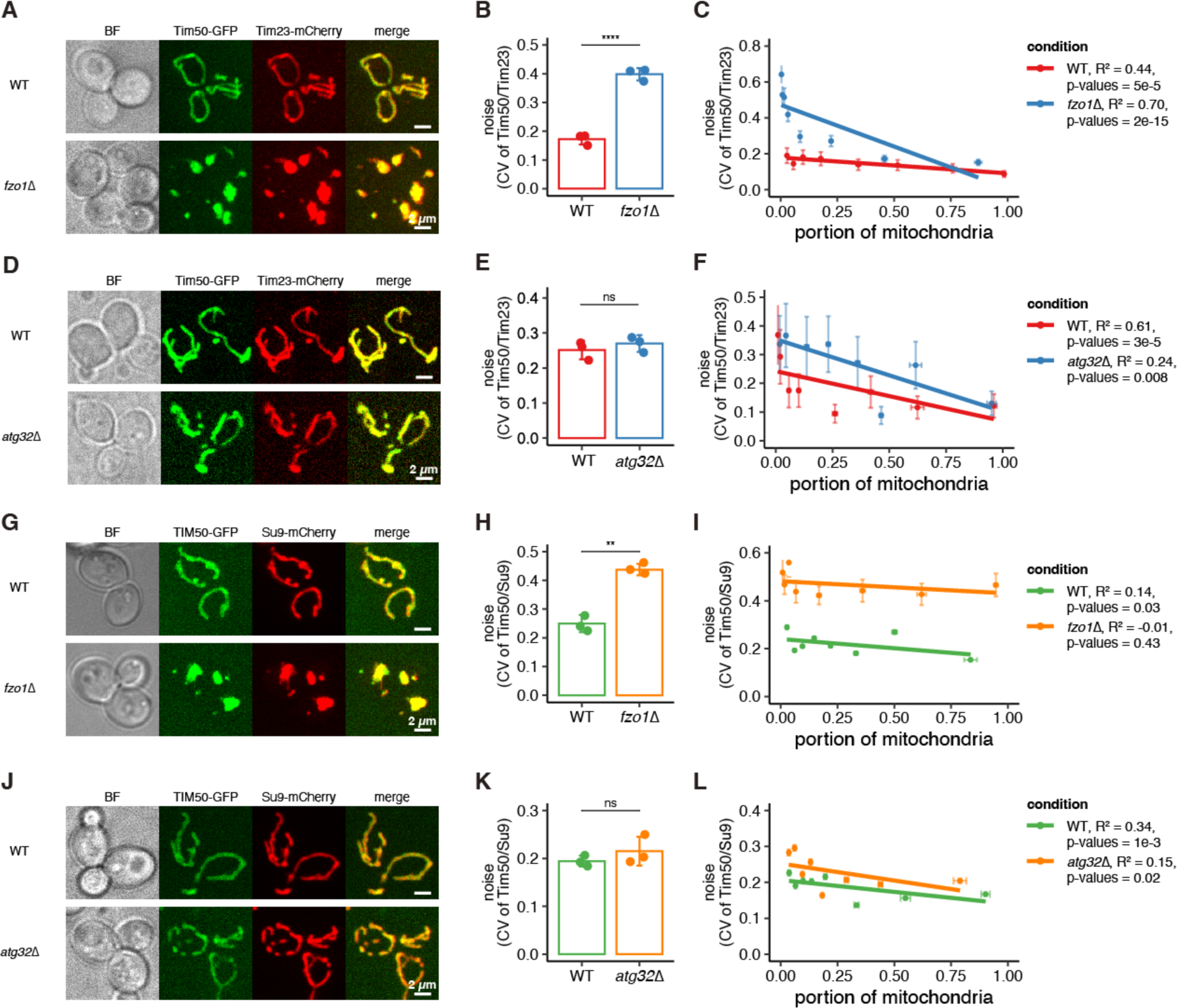
The heterogeneity of protein distribution is maintained by mitochondrial fission-fusion reaction and not through the mitophagy pathway. Protein distribution to each mitochondrial fragment in mutant strains: fission-mutant strain *fzo1*Δ (A-C, G-I), mitophagy-deficient strain *atg32*Δ (D-F, J-L). (A, D, G, J) The fluorescence of Tim50-GFP and Tim23-mCherry (A, D) or Su9-mCherry (G, J) were visualized by a spinning disc confocal microscope. BF: Bright field (center focal plane), Tim50-GFP, Tim23-mCherry, Su9-mCherry: z-projected images of each channel (GFP, RFP). Merge: merge of GFP and RFP channels. Scale bar, 2 µm. (B, E, H, K) The noise of protein distribution in mutant strains. The results represent the mean ± standard deviation of three independent experiments. Statistical significance was evaluated by Student t-test (**** P < 0.0001, ** P < 0.01). (C, F, I, L) Relationship between the noise of protein distribution and mitochondrial size in mutant strains. The noise value was calculated by dividing the population of a portion of mitochondria into eight groups along the portion of mitochondria. Error bars represent SEM of three independent experiments. R squared values and p-values for the F statistic hypothesis test are shown on the side.

### Inhibition of co-translational protein targeting prevents heterogeneous protein distribution

We next tested two possible strategies to reduce heterogeneity in the protein distribution for smaller mitochondrial fragments. Our computational modeling showed that increasing the number of mRNA copies would reduce heterogeneity. To test this, we introduced a GFP-tagged *TIM50* integration plasmid with different copy numbers to control mRNA copy numbers. We observed that an increase in copy number resulted in a decrease in noise in the *fzo1*Δ cells, indicating reduced heterogeneity in mitochondrial protein composition (Figure 4A). Doubling the mRNA copy number decreased noise by a factor of approximately 1.3 and tripling the copy number by approximately 1.8, similar to decreases predicted by modeling the mRNA production contribution to noise of 1.4 and 1.7, respectively. We further noticed that with an augmented expression of *TIM50* mRNA, there is a significant decrease in protein heterogeneity across all mitochondrial proportions (Figure 4B), which was also seen in our simulations (Figure 2F). We then examined if altering co-translational delivery by directing localization of mitochondrially localized mRNA to other specific subcellular locations affects the heterogeneity of the protein distribution. We tested a tethering model where a CAAX-tagged MCP protein (MCP-CAAX) represented plasma membrane tethering (Figure 4C) ^33^. In a WT strain, no significant effect of the plasma membrane tethering was observed compared to the control strain (Figure 4D). However, as consistent with the other experiments, heterogeneity in protein composition decreased with an increase in mitochondrial proportion across the strains. Although the noise level averaged over mitochondrial portion sizes is unaffected by CAAX tethering, we observed slightly reduced patterns of linear regression relationship in plasma membrane tethering (Figure 4E). Since we previously observed that *fzo1*Δ strains lost the regulation of protein heterogeneity with an increase in mitochondrial proportion, we analyzed the effect of membrane tethering on protein heterogeneity in the *fzo1*Δ cells. We found that membrane tethering successfully rescued the phenotype, and protein heterogeneity was significantly decreased compared to the control strain (Figure 4F, G). Collectively, the results suggest that mitochondrial protein heterogeneity can be inhibited by the increase in transcriptional level and a decrease in the co-translational protein delivery.

**Figure 4.**
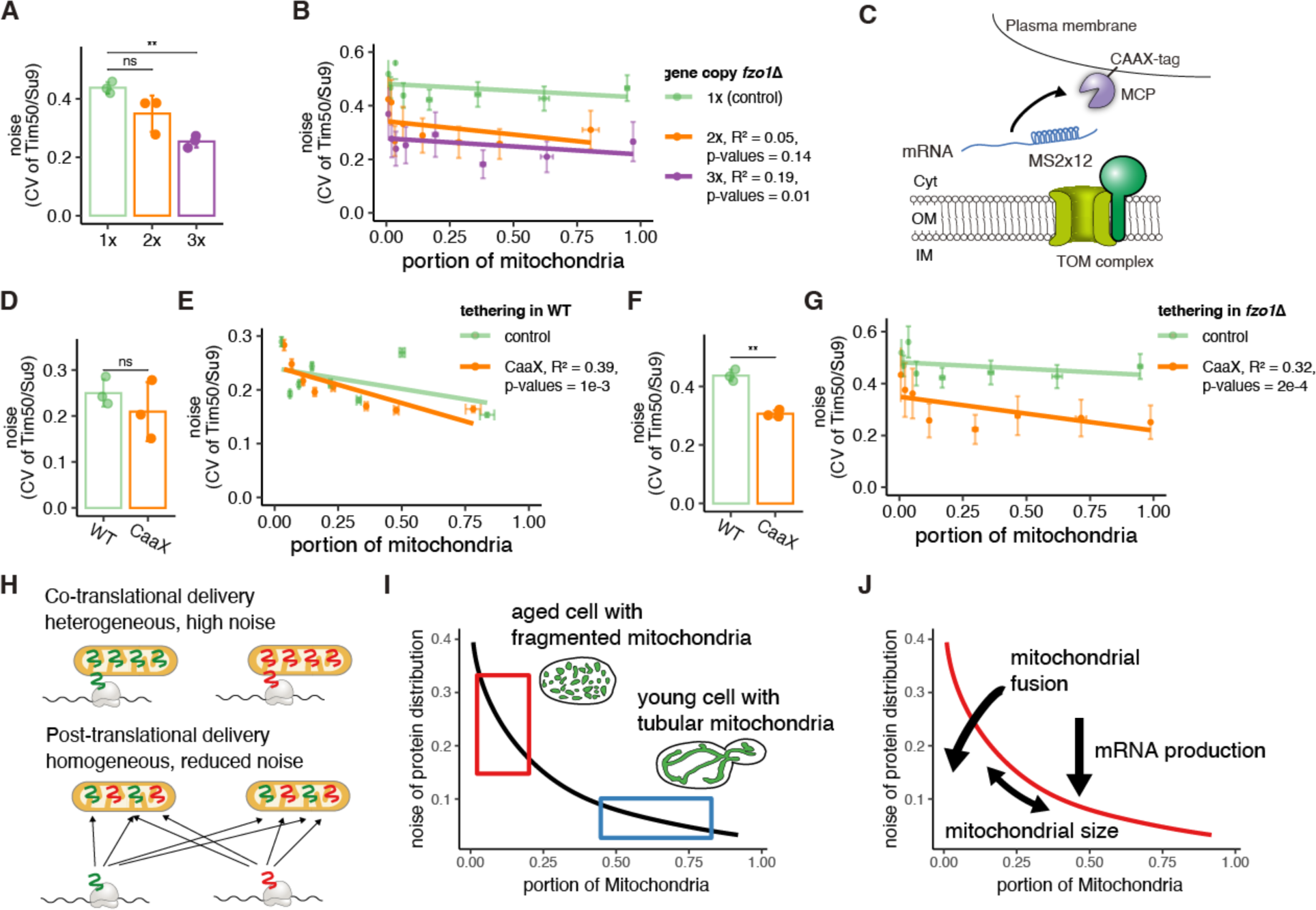
Co-translational protein delivery regulates protein composition. (A) The noise of protein distribution with variable copy numbers (1x, 2x, 3x) of GFP-tagged Tim50 integration plasmid in *fzo1*Δ strain (n>50). The results represent the mean ± standard deviation of three independent experiments. Statistical significance was evaluated by Student t-test (** P < 0.01). (B) Relationship between the noise of protein distribution and mitochondrial size in (A). The noise value was calculated by dividing the population of a portion of mitochondria into eight groups along the portion of mitochondria. Error bars represent SEM of three independent experiments. R squared values and p-values for the F statistic hypothesis test are shown on the side. (C) Schematic representation of plasma membrane tethering models. In plasma membrane tethering, MCP is bound to CAAX (a ras protein family member). (D, F) The noise of protein distribution in plasma membrane tethering (CAAX) of GFP-tagged Tim50 integration plasmid in WT (D) and *fzo1*Δ strain (F) (n>50). The results represent the mean ± standard deviation of three independent experiments. Statistical significance was evaluated by Student t-test (** P < 0.01). (E, G) Relationship between the noise of protein distribution and mitochondrial size in (D, F). The noise value was calculated by dividing the population of a portion of mitochondria into eight groups along the portion of mitochondria. Error bars represent SEM of three independent experiments. R squared values and p-values for the F statistic hypothesis test are shown on the side. (H) A schematic model description of different protein delivery mechanisms. Co-translational protein delivery increases protein distribution heterogeneity, while post-translational delivery reduces the noise in protein distribution. (I) Aged cells with fragmented mitochondria show higher noise, while young cells with tubular network mitochondrial structure show lower noise. (J) A schematic model description of mitochondrial protein distribution. Upon co-translational protein delivery, mitochondrial size influences stochastic mRNA localization. An increase in mitochondrial fission-fusion rates or mRNA production rates will each independently reduce protein distribution noise.

## Discussion

Aging is associated with an increase in mitochondrial fragmentation, as well as a decline in mitochondrial function across many species. Here, we report that the physical changes associated with mitochondrial fragmentation led to heterogeneity in protein distribution due to mRNA localization and co-translational protein delivery (Figure 4H, I). Our two-color fluorescent reporter gene analysis based on quantitative microscopy shows that mitochondrial proteins, even components of the same complex, are heterogeneously distributed to each mitochondrion in aged cells (Figure 1). There is an increase in heterogeneity as mitochondrial fragments become smaller. We observed that single-molecule *TIM50* mRNAs, which encode a co-translationally imported protein, remain stably associated with mitochondria, which suggests co-translational protein import may restrict protein production to specific mitochondrial fragments (Figure 2A-C). Using computational modeling, we found that the generation of heterogeneity can largely be explained by the stochastic nature of compartmentalization, and co-translational protein delivery is the main contributor to the high heterogeneity (Figure 2D-H, 4J). The heterogeneity of protein distribution was increased in mitochondrial fusion deficient *fzo1*Δ but not in the mitophagy deficient *atg32*Δ (Figure 3). This indicates that cells repress heterogeneity of protein distribution mainly by maintaining mitochondrial fission-fusion reactions and not through the mitophagy pathway. Lastly, we showed ways to reduce the noise level by either increasing the transcriptional level or decreasing co-translational protein delivery (Figure 4A-F).

We generally see highly complex mitochondrial networks maintained with mitochondrial fission and fusion dynamics across various cell types and conditions ^3^. We showed that a large and dynamic mitochondrial network structure, effectively forming a connected compartment, reduces noise levels by mixing the components across the mitochondrial network (Figure 1G, 3C, F, I, L, 4B, E, G). In particular, we found that mitochondrial fission and fusion reactions are essential to reduce mitochondrial protein concentration noise generated by mitochondrial fragmentation and co-translational protein delivery (Figure 3B, H). Our results suggest that when proteins do not form a complex, the heterogeneity in a single mitochondrial fragment can be substantially higher than when the proteins are part of the same complex (Figure 3C, I). This suggests that there may be pathways that inhibit protein heterogeneity of proteins forming complexes, such as protein degradation of excess subunits ^48^. While this study did not show a significant influence of the mitophagy pathway, the heterogeneity of the protein distribution throughout mitochondria (Figure 3E, K), regulation of protein production, and protein degradation may also contribute to more exact stoichiometries ^49,50^.

Early observations of inducible fragmentation due to loss of Fzo1 function found that while long-term fragmentation leads to loss of mtDNA and membrane potential, these phenotypes are not apparent immediately after fragmentation ^51^. This suggests that structural fragmentation alone does not drive mitochondrial dysfunction, but that mitochondrial dysfunction is caused by processes that occur over longer periods. As we have shown here, this nuclear-encoded mRNA noise-driven heterogeneity is a new and distinct proposal from stochasticity in mtDNA or mitophagy controlling heterogeneity and downstream dysfunction. Our observation indicates that co-translational protein targeting plays a significant role in generating protein heterogeneity. Even a slight initiation of mitochondrial fragmentation can have noticeable consequences due to the limited number of mRNA copy numbers compared to mtDNAs. From ∼25% to over 50% of nuclear-encoded mitochondrial mRNAs are localized to the mitochondria across both fermentative and respiratory conditions ^27,32,52^. With >100 mRNAs being localized even in fermentative conditions, the associated heterogeneity in protein expression due to co-translational insertion could be a strong driver of mitochondrial dysfunction as individual mitochondria deviate from the optimal stoichiometry of mitochondrial protein complexes. These postulate that overall mitochondrial dysfunction is introduced not only by mtDNA and mitophagy but also by co-translational protein targeting, which accelerate the dysfunction. Research is underway to better describe how mitochondrial fragmentation and co-translational protein import impact mitochondrial proteostasis. A finding of inhibitor compounds or genes in co-translational protein import is essential for this research direction. As many diseases and aging are associated with mitochondrial dysfunction, it is also important to determine whether there is a similar association between the degree of mitochondrial fragmentation and mitochondrial protein heterogeneity. Further studies focusing on this molecular mechanism can potentially contribute to the treatment of mitochondria-related diseases.

---

**Supplementary Figure 1.**
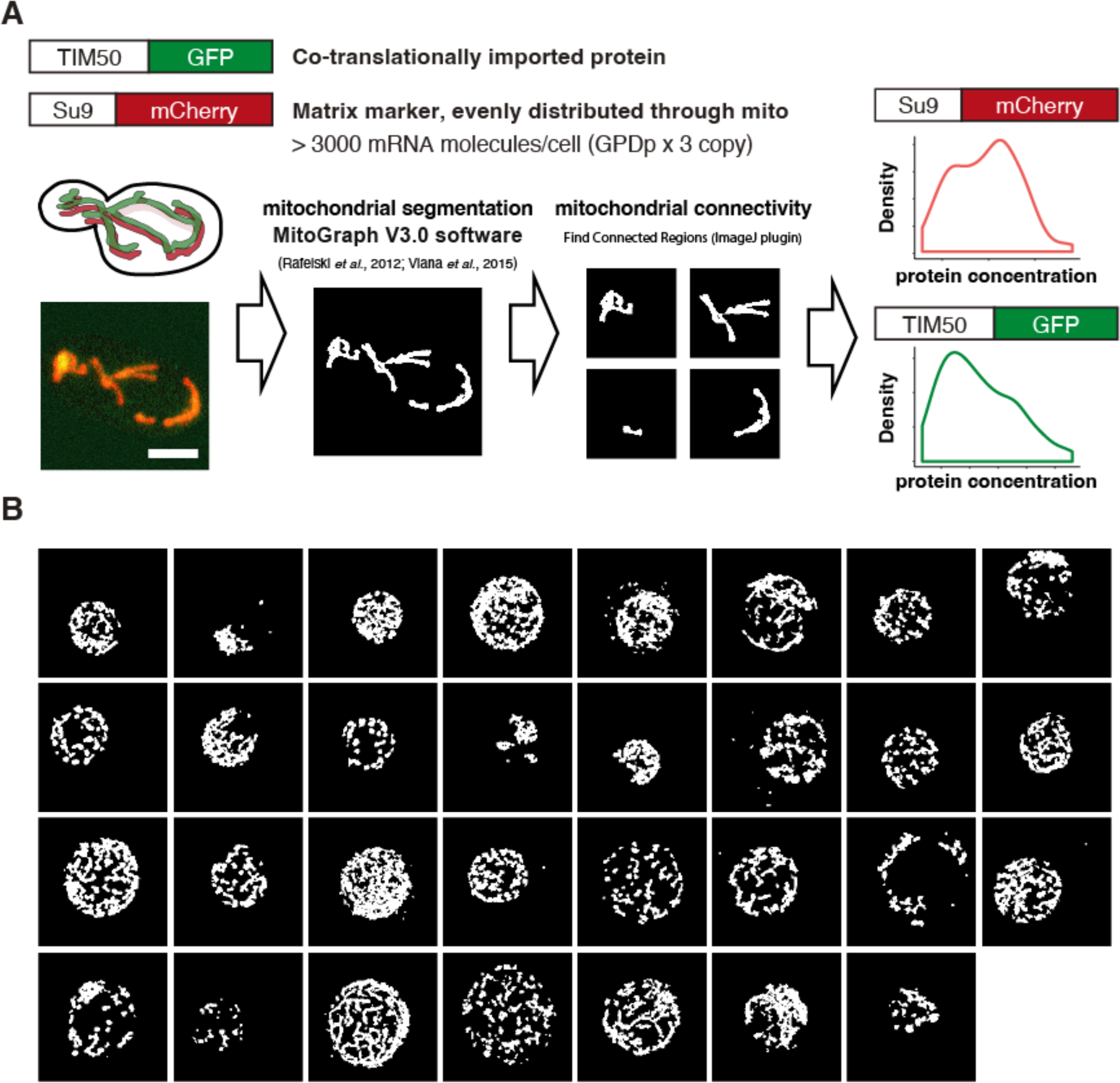
Quantification of mitochondrial protein composition. (A) To quantify the composition of mitochondrial proteins, we generated strains that express GFP conjugated to mitochondrial inner transporter protein Tim50 and mCherry conjugated to mitochondrial inner transporter protein Tim23 or mitochondrial matrix marker protein Su9. Su9-mCherry was expressed from the *TDH3* promoter of 3 copies of endogenously integrated plasmids. Tim50-GFP and Tim23-mCherry were expressed by the *TIM50* and *TIM23* promoters, respectively. Mitochondrial morphology was segmented from the RFP channel of the data using “MitoGraph V3.0 software”. We identified single fragments of mitochondria by analyzing the binary file of segmented mitochondria with the ImageJ plugin “Find Connected Regions.” The relative protein concentration of each fragment was measured based on the sum of each fragment’s intensity. Scale bar, 2 µm. (B) The cells used as aged cells 24 hours after the addition of estradiol. Z-projected segmented image of the RFP channel (Tim23-mCherry).

**Supplementary File 1**

**Yeast strains and plasmids used in this study**

**Supplementary File 2**

**List of oligonucleotides used for plasmid construction**

**Supplementary File 3**

**Row data of fluorescent intensity and size of mitochondrial fragments**

## Acknowledgments

We thank members of the Tsuboi laboratory and Zid laboratory for helpful discussions and feedback on the paper. This work was supported in part by startup funds from UCI, NSF grant MCB-1330451 and Ellison Medical Foundation (to S.M.R), startup funds from UCSD and the National Institutes of Health R35GM128798 (to B.M.Z.), and NINDS P30NS047101 (to UCSD microscopy Core). A.I.B. acknowledges support from a Natural Sciences and Engineering Research Council of Canada (NSERC) Discovery Grant and start-up funds provided by the Toronto Metropolitan University Faculty of Science. T.T. acknowledges support from the Japan Society for the Promotion of Science (JSPS) for a research abroad fellowship and postdoctoral fellowship (18J00995), Uehara Memorial Foundation for a research abroad fellowship, the Jilin Fuyuan Guan Food Group Co., Ltd, the Science, Technology, Innovation Commission of Shenzhen Municipality (WDZC20220811144737001), and startup funds from Tsinghua SIGS.

## Data and materials availability

The code used for analyzing the intensity of mitochondrial fragments is available from https://github.com/khan-ah. Further information and requests for resources, scripts, and reagents should be directed to and will be fulfilled by the lead contact, T.T.

## Methods

### Yeast strains and plasmids

The yeast strains and plasmids used are listed in Supplementary File 1, and the oligonucleotides used for plasmid construction and gene modification are listed in Supplementary File 2. To reduce variability among the constructed yeast strains, the strains were created either through the integration of a linear PCR product or a plasmid linearized through restriction digest. Tim50-GFP was expressed by *TIM50* promoter from integrated plasmid TTP155. The copy number of integrated plasmids in a cell was tested through microscopy screening, and the strains with one copy of integration were used for all the experiments except Figure 4. Fluorescent protein tagging for *TIM23* was performed with PCR-mediated homologous recombination using pFA6a-link-yomCherry-SpHis5 ^53^ and integrations were confirmed by PCR.

Su9-mCherry was expressed by GPD promoter from integrated plasmid TTP076, and the strains with three copies of integrated plasmids in a cell were screened through microscopy and used for further experiments. Deletion mutant strains were constructed with PCR-mediated homologous recombination using pFA6a-hphMX6 ^54^, and integrations were confirmed by PCR. Plasma membrane anchor MCP-iRFP-CaaX (TTP223) was constructed by swapping the GFP of TTP167 with the iRFP of TTP145 through the combination of Gibson assembly and PCR ^33^. The strains expressing MCP-iRFP-CaaX from two sets of integrated plasmids were selected through microscopy screening and used for further experiments.

### Microscopy

Image data for C-terminal integrated fluorescent proteins were collected as follows: Yeast cells were grown in the YPAD (YPA medium containing 2% glucose) with a 15ml glass tube at 30℃ with rotator speeds of 60 rpm. 100μl of mid-log phase wild-type yeast cells (OD600 of 0.4 to 0.7) were harvested and placed into a 96-well Glass Bottom Plate (Cellvis LLC) coated with 0.1 mg/mL concanavalin A (Sigma-Aldrich C2010). For the aging experiment, we used mother enrichment program (MEP) strains, which were controlled by the addition of estradiol (final l μM) (Sigma-Aldrich E2758). Cells were imaged at 23℃ with 3-second intervals by an Eclipse Ti2-E Spinning Disk Confocal with Yokogawa CSU-X1 (Yokogawa) with 50 µm pinholes, located at the Nikon Imaging Center UCSD. Imaging was performed using SR HP APO TIRF 100x 1.49 NA oil objective with the correction collar set manually for each experiment (pixel size 0.0936µm). Z-stacks (200nm steps) were acquired by a Prime 95B sCMOS camera (Photometrics). Imaging was controlled using NIS-Elements software (Nikon). Image data for Figure 2D-F were imaged at 23℃ by an Eclipse Ti2-E Spinning Disk Confocal with Yokogawa CSU-W1 (Yokogawa) with 50 µm pinholes, located at the Tsinghua-SIGS Tsuboi Laboratory. Imaging was performed using CFI Plan Apochromat Lambda 100x 1.49 NA oil objective. Z-stacks (200nm steps) were acquired by a Prime 95B sCMOS camera (Photometrics) (pixel size 0.11µm). Imaging was controlled using NIS-Elements software (Nikon). Single-molecule mRNA visualization with mitochondria was performed as follows: Yeast cells were grown in the YPAD with a 15ml glass tube at 30℃ with rotator speeds of 60 rpm. 300μl of mid-log phase wild-type yeast cells (OD600 of 0.4-0.7), grown in an appropriate medium, were harvested and placed into a Y04C microfluidic chamber controlled by the CellASIC Onix system. 300μl YPAD were placed into the flow-wells, the chambers were loaded with cells at 3psi, and the medium continuously flowed at 3psi. Cells were imaged at 30℃ with a Yokogawa CSU-X1 Spinning Disk Confocal (Solamere Technology Group) mounted on a Nikon Eclipse Ti chassis motorized inverted microscope, located at the Department of Developmental & Cell Biology UCI. Imaging was performed using a 100x/1.49 NA oil APO TIRF objective with the correction collar set manually for each experiment and a 1x tube lens (pixel size 0.084μm). Z-stacks (300nm steps) were acquired in the fluorescent channel (33ms exposure) on a Hamamatsu electron-multiplying charge-coupled device (EMCCD) camera. Imaging was controlled using MicroManager ImageAcquisition (v1.4.16). The experiments for Figure 3G-I and 4A-G were conducted on the same day.

### Quantification of protein expression in each mitochondrion from image data

Quantification of GFP-tagged mitochondrially localized protein expression was performed using a custom analysis pipeline. The MitoGraph was applied to the mCherry channel to identify the locations of mitochondria, and the resulting binary mask was then used to segment the GFP channels of individual mitochondria. Next, the sum of the intensity of the GFP channel covered by a binary mask from each mitochondrion was measured using a custom ImageJ script.

### Reconstruction of 3D mitochondria and mRNA visualization

To allow accurate visualization of mRNA molecules, multiple MS2 stem-loops are inserted in the 3’-UTR of the mRNA of interest and are recognized by the MCP-GFP fusion protein ^55,56^. We improved this system by titrating the MCP-GFP levels until we observed single-molecule mRNA foci ^33^. We then performed rapid 3D live cell imaging using spinning disk confocal microscopy. We reconstructed and analyzed the spatial relationship between the mRNAs and mitochondria using the custom ImageJ plugin Trackmate ^57^ and MitoGraph V2.0, which we previously developed to reconstruct 3D mitochondria based on matrix marker fluorescent protein intensity ^33,58,59^. We measured the distance between mRNA and mitochondria by finding the closest meshed surface area of the mitochondria matrix ^33^. The code used for the analysis is available from https://github.com/tsuboitat.

### mRNA production and protein delivery noise modeling

If mRNA is produced at a constant rate, the number of mRNA produced over a generation time period will be Poisson-distributed. With a *TIM50* mRNA copy number of approximately six ^33^ and a lifetime of approximately 10 minutes ^45^, there will be a mean of approximately 54 mRNA produced in a 90 minutes generation. Poisson-distributed processes have an equal mean and variance, such that the coefficient of variation is the inverse square root of the mean of 54 or a coefficient of variation of 0.14 for isolated mRNA production noise.

For isolated protein delivery noise, we consider proteins that are delivered to mitochondria in N_del_ delivery events, with each mitochondrion selected with a probability proportional to the mitochondrial size. With the mitochondrial size that is a fraction f of the total mitochondrial size in the cell, then for Poisson-distributed delivery events, a mitochondrion will have a mean and variance of fN_del_ delivery events, and thus a fractional size f-dependent coefficient of variation of (fN_del_)^-1/2^. If each mRNA docks at one mitochondrion and delivers all its translated proteins to that mitochondrion, then the size-dependent coefficient of variation over a generation is (fN_del_)^-1/2^ = (54f)^-1/2^. For mRNA that does not dock at a mitochondrion, each translated protein stochastically selects a mitochondrion proportional to its size, with the copy number of 4000 Tim50 proteins ^26^, each generation the coefficient of variation is (fN_del_)^-1/2^ = (4000f)^-1/2^.

### In silico Experiment (Stochastic simulation)

mRNA production and decay, protein production and decay, mitochondrial fusion, and cell division are all stochastically simulated for two mitochondria, one of fractional size f and the other with fractional size 1 – f, with the Gillespie algorithm^60,61^. Two mitochondria are the mean number of connected components in yeast mitochondrial networks ^62^. With *TIM50* mRNA copy number of approximately 5 ^33^ and mRNA lifetime of approximately 10 minutes ^45^, then mRNA is produced at the rate of 5/lifetime, and each mRNA decays at the rate of 1/lifetime. Proteins are produced at a rate of 0.13/s per *TIM50* mRNA, corresponding to the rate at which 4000 proteins will typically be present immediately prior to division and similar to an earlier estimate of the rate of *TIM50* translation initiation of 0.126/s ^63^. Each protein decays at a rate of 1/7200 per second, corresponding to a 2 hour lifetime ^64^. For the model of co-translational protein delivery to mitochondria, mRNA selects a mitochondrion when produced and delivers all proteins to that mitochondrion. For post-translational protein delivery to mitochondria, a mitochondrion for the protein to be delivered is selected for each protein produced. Mitochondrial fusion is represented in the model by events that equalize mitochondrial protein concentrations across mitochondria, occurring at a certain rate. We explore fusion event rates of zero and the range of 10^-3.5^ – 10^-2.5^/s, similar to experimentally estimated mitochondrial fusion rates ^24,65,66^. Cell division is implemented by instantly halving the protein numbers in each mitochondrion every generation period of 90 minutes. Data shown in Figures 2E-G is after simulating for ten generations.

## Supplemental information

Supplemental Information includes 1 figure and 3 files.

## References

1. Spinelli, J. B. & Haigis, M. C. The multifaceted contributions of mitochondria to cellular metabolism. Nat Cell Biol 20, 745–754 (2018).

2. Eisner, V., Picard, M. & Hajnóczky, G. Mitochondrial dynamics in adaptive and maladaptive cellular stress responses. Nat Cell Biol 20, 755–765 (2018).

3. Westermann, B. Mitochondrial fusion and fission in cell life and death. Nature Reviews Molecular Cell Biology vol. 11 872–884 (Nature Publishing Group, 2010).

4. das Neves, R. P., et al. Connecting variability in global transcription rate to mitochondrial variability. PLoS Biol 8, (2010).

5. Johnston, I. G. et al. Mitochondrial variability as a source of extrinsic cellular noise. PLoS Comput Biol 8, 35–37 (2012).

6. El-Hattab, A. W. & Scaglia, F. Mitochondrial DNA Depletion Syndromes: Review and Updates of Genetic Basis, Manifestations, and Therapeutic Options. Neurotherapeutics 2013 10:2 10, 186–198 (2013).

7. Aryaman, J., Johnston, I. G. & Jones, N. S. Mitochondrial heterogeneity. Front Genet 10, 1–16 (2019).

8. Chen, H. et al. Mitochondrial fusion is required for mtDNA stability in skeletal muscle and tolerance of mtDNA mutations. Cell 141, (2010).

9. Johnston, I. G. & Jones, N. S. Closed-form stochastic solutions for non-equilibrium dynamics and inheritance of cellular components over many cell divisions. Proceedings of the Royal Society A: Mathematical, Physical and Engineering Sciences 471, (2015).

10. Stephan, T. et al. MICOS assembly controls mitochondrial inner membrane remodeling and crista junction redistribution to mediate cristae formation. EMBO J 39, 1–24 (2020).

11. Jimenez, L., Laporte, D., Duvezin-Caubet, S., Courtout, F. & Sagot, I. Mitochondrial ATP synthases cluster as discrete domains that reorganize with the cellular demand for oxidative phosphorylation. J Cell Sci 127, 719–726 (2014).

12. Gerencser, A. A. et al. Quantitative measurement of mitochondrial membrane potential in cultured cells: calcium-induced de- and hyperpolarization of neuronal mitochondria. J Physiol 590, 2845– 2871 (2012).

13. Gerencser, A. A., Mookerjee, S. A., Jastroch, M. & Brand, M. D. Measurement of the Absolute Magnitude and Time Courses of Mitochondrial Membrane Potential in Primary and Clonal Pancreatic Beta-Cells. PLoS One 11, e0159199 (2016).

14. Wang, W. et al. Superoxide Flashes in Single Mitochondria. Cell 134, 279–290 (2008).

15. Santo-Domingo, J., Giacomello, M., Poburko, D., Scorrano, L. & Demaurex, N. OPA1 promotes pH flashes that spread between contiguous mitochondria without matrix protein exchange. EMBO J 32, 1927–1940 (2013).

16. Yoon, Y., Galloway, C. A., Jhun, B. S. & Yu, T. Mitochondrial dynamics in diabetes. Antioxid Redox Signal 14, 439–457 (2011).

17. Galloway, C. A. & Yoon, Y. Mitochondrial morphology in metabolic diseases. Antioxid Redox Signal 19, 415–430 (2013).

18. Gao, J. et al. Abnormalities of Mitochondrial Dynamics in Neurodegenerative Diseases. (2017) doi:10.3390/antiox6020025.

19. Liu, Y. J., McIntyre, R. L., Janssens, G. E. & Houtkooper, R. H. Mitochondrial fission and fusion: A dynamic role in aging and potential target for age-related disease. Mech Ageing Dev 186, (2020).

20. Zhao, J. et al. Mitochondrial dynamics regulates migration and invasion of breast cancer cells. Oncogene 32, 4814–4824 (2013).

21. Xu, K. et al. MFN2 suppresses cancer progression through inhibition of mTORC2/Akt signaling. Sci Rep 7, (2017).

22. Li, Y. et al. A programmable fate decision landscape underlies single-cell aging in yeast. Science (1979) (2020) doi:10.1126/science.aax9552.

23. Hughes, A. L. & Gottschling, D. E. An early age increase in vacuolar pH limits mitochondrial function and lifespan in yeast. Nature 492, 261–265 (2012).

24. Twig, G. et al. Fission and selective fusion govern mitochondrial segregation and elimination by autophagy. EMBO J 27, 433–446 (2008).

25. Hoitzing, H., Johnston, I. G. & Jones, N. S. What is the function of mitochondrial networks? A theoretical assessment of hypotheses and proposal for future research. BioEssays 37, 687–700 (2015).

26. Morgenstern, M. et al. Definition of a High-Confidence Mitochondrial Proteome at Quantitative Scale. Cell Rep 19, 2836–2852 (2017).

27. Marc, P. et al. Genome-wide analysis of mRNAs targeted to yeast mitochondria. EMBO Rep 3, 159–64 (2002).

28. Saint-Georges, Y. et al. Yeast mitochondrial biogenesis: a role for the PUF RNA-binding protein Puf3p in mRNA localization. PLoS One 3, e2293 (2008).

29. Gadir, N., Haim-Vilmovsky, L., Kraut-Cohen, J. & Gerst, J. E. Localization of mRNAs coding for mitochondrial proteins in the yeast Saccharomyces cerevisiae. RNA 17, 1551–65 (2011).

30. Garcia, M. et al. Mitochondria-associated yeast mRNAs and the biogenesis of molecular complexes. Mol Biol Cell 18, 362–368 (2007).

31. Fazal, F. M. et al. Atlas of Subcellular RNA Localization Revealed by APEX-Seq. Cell 178, 473–490.e26 (2019).

32. Williams, C. C., Jan, C. H. & Weissman, J. S. Targeting and plasticity of mitochondrial proteins revealed by proximity-specific ribosome profiling. Science (1979) 346, 748–751 (2014).

33. Tsuboi, T. et al. Mitochondrial volume fraction and translation speed impact mRNA localization and production of nuclear-encoded mitochondrial proteins. Elife 9, 1–24 (2020).

34. Eliyahu, E. et al. Tom20 Mediates Localization of mRNAs to Mitochondria in a Translation-Dependent Manner. Mol Cell Biol 30, 284–294 (2010).

35. Lesnik, C., Cohen, Y., Atir-Lande, A., Schuldiner, M. & Arava, Y. OM14 is a mitochondrial receptor for cytosolic ribosomes that supports co-translational import into mitochondria. Nat Commun 5, 1–10 (2014).

36. Borst, P. & Grivell, L. A. The Mitochondrial Genome of Yeast. Cell 15, 705–723 (1978).

37. Palmer, C. S., Anderson, A. J. & Stojanovski, D. Mitochondrial protein import dysfunction: mitochondrial disease, neurodegenerative disease and cancer. FEBS Letters vol. 595 Preprint at 10.1002/1873-3468.14022 (2021).

38. Demishtein-Zohary, K. & Azem, A. The TIM23 mitochondrial protein import complex: function and dysfunction. Cell and Tissue Research vol. 367 Preprint at 10.1007/s00441-016-2486-7 (2017).

39. Lindstrom, D. L. & Gottschling, D. E. The mother enrichment program: A genetic system for facile replicative life span analysis in Saccharomyces cerevisiae. Genetics 183, 413–422 (2009).

40. Brennan, C. M. et al. Protein aggregation mediates stoichiometry of protein complexes in aneuploid cells. Genes Dev 33, (2019).

41. Miura, F. et al. Absolute quantification of the budding yeast transcriptome by means of competitive PCR between genomic and complementary DNAs. BMC Genomics 9, 1–14 (2008).

42. Lahtvee, P. J. et al. Absolute Quantification of Protein and mRNA Abundances Demonstrate Variability in Gene-Specific Translation Efficiency in Yeast. Cell Syst 4, (2017).

43. Marguerat, S. et al. Quantitative analysis of fission yeast transcriptomes and proteomes in proliferating and quiescent cells. Cell 151, (2012).

44. Munchel, S. E., Shultzaberger, R. K., Takizawa, N. & Weis, K. Dynamic profiling of mRNA turnover reveals gene-specific and system-wide regulation of mRNA decay. Mol Biol Cell 22, 2787–2795 (2011).

45. Chan, L. Y., Mugler, C. F., Heinrich, S., Vallotton, P. & Weis, K. Non-invasive measurement of mRNA decay reveals translation initiation as the major determinant of mRNA stability. Elife 7, 1– 32 (2018).

46. Onishi, M., Yamano, K., Sato, M., Matsuda, N. & Okamoto, K. Molecular mechanisms and physiological functions of mitophagy. EMBO J 40, (2021).

47. Schmidt, B., Hennig, B., Zimmermann, R. & Neupert, W. Biosynthetic pathway of mitochondrial ATPase subunit 9 in Neurospora crassa. Journal of Cell Biology 96, (1983).

48. McShane, E. et al. Kinetic Analysis of Protein Stability Reveals Age-Dependent Degradation. Cell 167, (2016).

49. Song, J., Herrmann, J. M. & Becker, T. Quality control of the mitochondrial proteome. Nature Reviews Molecular Cell Biology vol. 22 Preprint at 10.1038/s41580-020-00300-2 (2021).

50. Couvillion, M. T., Soto, I. C., Shipkovenska, G. & Churchman, L. S. Synchronized mitochondrial and cytosolic translation programs. Nature 533, 499–503 (2016).

51. Hermann, G. J. et al. Mitochondrial fusion in yeast requires the transmembrane GTPase Fzo1p. Journal of Cell Biology 143, (1998).

52. Tsuboi, T., Leff, J. & Zid, B. M. Post-transcriptional control of mitochondrial protein composition in changing environmental conditions. Biochem Soc Trans 48, 2565–2578 (2020).

53. Lee, S., Lim, W. A. & Thorn, K. S. Improved Blue, Green, and Red Fluorescent Protein Tagging Vectors for S. cerevisiae. PLoS One 8, 4–11 (2013).

54. Goldstein, A. L. & McCusker, J. H. Three new dominant drug resistance cassettes for gene disruption in Saccharomyces cerevisiae. Yeast (1999) doi:10.1002/(SICI)1097-0061(199910)15:14<1541::AID-YEA476>3.0.CO;2-K.

55. Bertrand, E. et al. Localization of ASH1 mRNA Particles in Living Yeast. Mol Cell 2, 437–445 (1998).

56. Haim-Vilmovsky, L. & Gerst, J. E. m-TAG: a PCR-based genomic integration method to visualize the localization of specific endogenous mRNAs in vivo in yeast. Nat Protoc 4, 1274–1284 (2009).

57. Tinevez, J. Y. et al. TrackMate: An open and extensible platform for single-particle tracking. Methods (2017) doi:10.1016/j.ymeth.2016.09.016.

58. Rafelski, S. M. et al. Mitochondrial network size scaling in budding yeast. Science 338, 822–4 (2012).

59. Viana, M. P., Lim, S. & Rafelski, S. M. Quantifying mitochondrial content in living cells. Methods in Cell Biology vol. 125 (Elsevier Ltd, 2015).

60. Gillespie, D. T. Stochastic simulation of chemical kinetics. Annu Rev Phys Chem 58, (2007).

61. Gillespie, D. T. Exact stochastic simulation of coupled chemical reactions. in Journal of Physical Chemistry vol. 81 (1977).

62. Viana, M. P. et al. Mitochondrial Fission and Fusion Dynamics Generate Efficient, Robust, and Evenly Distributed Network Topologies in Budding Yeast Cells. Cell Syst 10, (2020).

63. Arceo, X. G., Koslover, E. F., Zid, B. M. & Brown, A. I. Mitochondrial mRNA localization is governed by translation kinetics and spatial transport. PLoS Comput Biol 18, (2022).

64. Martin-Perez, M. & Villén, J. Determinants and Regulation of Protein Turnover in Yeast. Cell Syst 5, (2017).

65. Sukhorukov, V. M., Dikov, D., Reichert, A. S. & Meyer-Hermann, M. Emergence of the Mitochondrial Reticulum from Fission and Fusion Dynamics. PLoS Comput Biol 8, (2012).

66. Patel, P. K., Shirihai, O. & Huang, K. C. Optimal Dynamics for Quality Control in Spatially Distributed Mitochondrial Networks. PLoS Comput Biol 9, (2013).

