## Supplementary_file_1 for "Mitochondrial protein heterogeneity stems from the stochastic nature of co-translational protein targeting in cell senescence"

**Supplementary File 1. Yeast strains and plasmids used in this study**

Strain/plasmids	Genotype[plasmid](plasmid number)	Source

Strains

W303-1a	MATa *ade2-1 can1-100 his3-11 leu2-3 trp1-1 ura3*	Lab stock

UCC8853 MATa *his3 leu2 lys2∆ ura3∆0 ho*∆::*PSCW11-cre-EBD78-NATMX*	Lindstrom and Gottschling, 2009
*loxP-CDC20-Intron-loxP-HPHMX loxP-UBC9-loxP-LEU2*

TTY1089	W303-1a TIM50::12xMS2tag::TIM50^3’UTR^, TTP076, TTP080	Tsuboi et al., 2020

W303-1a	MATa ade2-1 can1-100 his3-11 leu2-3 trp1-1 *fzo1*Δ::*hphMX*	This study

W303-1a	MATa ade2-1 can1-100 his3-11 leu2-3 trp1-1 *atg32*Δ::*hphMX*	This study

Plasmids
pFA6a-link-yomCherry-CaURA3	Lee et al., 2013

pFA6a-hphMX6	Goldstein and McCusker, 1999

TTP076	pRS406*GPD*p-Su9-mCherry	Tsuboi et al., 2020
TTP080	pRS405*CYC1*p-MS2-4xGFP	Tsuboi et al., 2020

TTP145	pRS403*TIM50p-TIM50mts(1-300)-TIM50cds-flagiRFP-TIM50ter-MS2tag*	Tsuboi et al., 2020

TTP155	pRS403*TIM50p-TIM50mts(1-300)-TIM50cds-flagyoGFP-TIM50ter-MS2tag*	Tsuboi et al., 2020

TTP167	pRS405*CYC1*p-MS2-2xGFP-CaaX	Tsuboi et al., 2020

TTP223	pRS405*CYC1*p-MS2-iRFP-CaaX	This study
