## Supplementary_file_2 for "Mitochondrial protein heterogeneity stems from the stochastic nature of co-translational protein targeting in cell senescence"

**Supplementary File 2. List of oligonucleotides used for the experiment**

Gene Name	Oligonucleotides used for plasmid construction

*TTO465_Fzo1D::MX4/6 F	TTATCTGATATCACGGATAGAGGCAAAACGGTAGGCTCATTTAACGcggatccccgggttaattaaggcg*

*TTO466_Fzo1D::MX4/6 R	ACATTATGTATATTGATTTGAAAAGACCTCATATATTTACAAGAATATcataggccactagtggatctga*

*TTO668_atg32D::MX4/6 F	GTCCTAATCACAAAAGCAAAAAAAATCTGCCAGGAACAGTAAACATcggatccccgggttaattaaggcg*

*TTO669_atg32D::MX4/6 R	GTAAAAAAGTGAGTAGGAACGTGTATGTTTGTGTATATTGGAAAAAGGcataggccactagtggatctga*

*TTO631_TIM23-Kurt S. Thorn F	TGCGCCGTCTGGTGTAGTGTCAAGAAAAGACTACTTGAAAAAggtgacggtgctggttta*

*TTO632_TIM23-Kurt S. Thorn R	AGAGAGAGAGAAAGAGAGAGAGAGAGTAGGTTCTTGTGTTGCtcgatgaattcgagctcg*

*TTO633_TIM44-Kurt S. Thorn F	AAGATCTTGGAGTTTGTGCGCGGGGGTTCTAGACAATTCACCggtgacggtgctggttta*

*TTO634_TIM44-Kurt S. Thorn R	GAAGGAAAAGGAAAAGAAAACAAAAGAGTACATCGAAACCAAtcgatgaattcgagctcg*

*TTO586_CaaX_F	ggtagcgatagcgcaggcagtgctg*

*TTO599_MCP_reverse-4	GATATCGAATTCCTGCAGCCCGGGGGATCCGTAGATGCCGGAGTTTGC*

*TTO589_GA_MCP_iRFP	ggctgcaggaattcgatatcgtgATGGACTACAAGGACGACGATGACAAG*

*TTO590_GA_iRFP_CaaX	cagcactgcctgcgctatcgctaccttcttccataacaccgatttgccaagc*
